# Predicting E. coli concentrations using limited qPCR deployments at Chicago beaches

**DOI:** 10.1101/250480

**Authors:** Nick Lucius, Kevin Rose, Callin Osborn, Matt E Sweeney, Renel Chesak, Daniel Y. Little, Scott Beslow, Tom Schenk

## Abstract

Culture-based methods to measure *Escherichia coli* (*E. coli*) are used by beach administrators to inform whether bacteria levels represent an elevated risk to swimmers. Since results take up to 12 hours, statistical models are used to forecast bacteria levels in lieu of test results; however they underestimate days with elevated fecal indicator bacteria levels. Quantitative polymerase chain reaction (qPCR) tests return results within 3 hours but are 2 to 5 times more expensive than culture-based methods. This paper presents a prediction model which uses limited deployments of qPCR tested sites with inter-beach correlation to predict when bacteria will exceed acceptable thresholds. The model can be used to inform management decisions on when to warn residents or close beaches due to exposure to the bacteria. Using data from Chicago collected between 2006 and 2016, the model proposed in this paper increased sensitivity from 3.4 percent to 11.2 percent–a 230 percent increase. We find that the correlation between beaches are substantial enough to provide higher levels of precision and sensitivity to predictive models. Thus, limited deployments of qPCR testing can be used to deliver better predictions for beach administrators at lower cost and less complexity.

## 1. Introduction

Swimming in recreational waters contaminated with high levels of fecal indicator bacteria (FIB) is associated with higher incidence of gastrointestinal (GI) illnesses (Prüss 1998). The risk of exposure is associated with the amount of FIB measured in the water. A logarithmic increase in average *Enterococcus* was associated with a 1.4 increase in the odds ratio of contracting GI illness (Wade et al. 2006). Evidence also suggests that children are more likely to contract GI illnesses when exposed to the same contaminated water as adults (Wade et al. 2008).

To prevent this, managers of recreational beaches use culture-based methods to measure FIB levels. Sampling is typically conducted early in the morning, but results take upward of 12 hours (Kinzelman et al. 2003). Between the time of sampling and subsequent results, beach conditions will often change so the water sample is not relevant to today’s beach conditions (Whitman, Nevers, and Gerovac 1999; Boehm et al. 2002). To get around this, researchers have developed statistical models–dubbed “nowcast” models–to estimate FIB for the day based on the previous day *E. coli* levels and factors such as precipitation, wind, and water conditions (Francy et al. 2013).

These models often incorrectly predict that beaches will not have elevated FIB levels–known as Type II errors or “false negatives” (Nevers and Whitman 2011; Rabinovici et al. 2004; Boehm et al. 2002). For instance in 2015, Chicago beaches had 200 events where FIB levels were too high; however, only 14 (7 percent) of these “beach days” were forecasted by an existing predictive model used by the Chicago Park District. These models do have good overall fit, but elevated FIB levels are statistically rare events and models often fail to predict them–known as low “sensitivity”.

Meanwhile, scientists have developed new methods which measure FIB levels in water with substantially less delay. *Enterococci* quantitative polymerase chain reaction (qPCR) methods can determine FIB levels within 3 to 4 hours and yield similar results as culture-based methods (Haugland et al. 2005; Kinzelman et al. 2003). The rapid turnaround lets managers determine warnings or closures based on tests within the same day instead of relying on statistical models.

However, this approach has a drawback of cost and equipment availability. qPCR testing can cost between 2 to 5 times more than traditional culture-based methods (Bienkowski 2017). Thus, managers are faced with a dilemma of choosing between expensive qPCR methods or choosing slower culture-based methods and using predictive models to produce swim advisories.

Both of these tests have thresholds for acceptable FIB levels. The Environmental Protection Agency (EPA) publishes water quality criteria in accordance with the Clean Water Act. The current EPA recreational water quality criteria accepts culture-based or qPCR-based methods (Environmental Protection Agency 2012). Acceptable levels for culture-based methods of testing *E. coli* should not exceed 235 CFU/100 ml while acceptable levels for qPCR testing of *enterococci* should not exceed 1,000 cell equivalents (CE) (Byappanahalli et al. 2010). These rules can be leveraged to create a new predictive model which mixes the short turn-around of qPCR with statistical models to produce swim advisories.

In this study, we exploit the historical correlation between beaches to estimate FIB readings. We also use limited *enterococci* qPCR results at specific beaches to predict elevated *E. coli* levels at other beaches using clustering algorithms and random forest regressions. This hybrid approach allows limited deployment of qPCR equipment to reduce overall costs and provides a higher quality statistical model.

We use 10 years of historical *E. coli* measurements from 20 beaches in Chicago to create a model to forecast whether *E. coli* levels at a beach will be elevated.

## 2. Material and Methods

### 2.1. Prior-day Nowcast Models

Timing is a crucial factor for monitoring FIB levels. We adopt the typical shorthand to denote time periods by *t* to denote now; *t* – 1, *t* – 2, *t* – *n* to denote last period, two periods ago, and *some* periods ago; and *t* + 1, *t* + 2, *t* + *n* to denote the next period, two periods from now, and *some* period in the future. For the purposes of this paper, we often use *t* to denote *today*, *t* – 1 to denote *yesterday*, and *t* +1 to denote *tomorrow*.

The defining characteristic of prior-day nowcast models is using FIB levels from the previous day (*t* – 1) to predict bacteria levels for the current day (*t*). Models will also incorporate covariates or predictors to improve the accuracy of models, such as precipitation (Ackerman and Weisberg 2003; Morrison et al. 2003), sunlight (Whitman et al. 2004), wind (Smith et al. 1999; Olyphant and Whitman 2004), wave and tidal levels (Le Fevre and Lewis 2003; Crowther, Kay, and Wyer 2001), lake levels (Francy, Bertke, and Darner 2009), turbidity (Olyphant et al. 2003), and density of humans and animals (Boehm et al. 2003; Reeves et al. 2004).

These models rely on prior-day data since culture-based testing is not available until upward of 12 hours after the samples were collected. The prediction at beach *x_i_* at time *t* is dependent on the prior periods culture-based results, 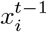 and aforementioned covariates, *w*_1_, *w*_2_, …, *w_j_*. These models often take the form of

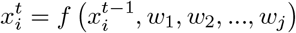

where *f* (…) is a some function or algorithm that inputs raw data and outputs a probability. For instance, the linear regression model is typically 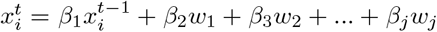, where *β_j_* are the coefficients that weight the importance of each input.

Various models are used to improve accuracy, such as log transformations (Nevers and Whitman 2005), polynomial coefficients (Frick, Ge, and Zepp 2008), logistic regression, partial least squares (Hou, Rabinovici, and Boehm 2006), generlized boosted regression modeling (Cyterski et al. 2014), random forest, and genetic algorithm approaches (Brooks et al. 2016). These classes of models often has the same structure of using FIB levels from time *t* ≠ 1–the prior-day–and covariates from time *t* to predict beaches at time *t*.

Yet, the reliance on prior-day FIB levels are likely the source of inaccuracy. The cause of elevated levels is unlikely to persist between days (Morrison et al. 2003; Cheung, Chang, and Hung 1991; Brenniman, Rosenberg, and Northrop 1981). Covariates can help determine when conditions are optimal for bacteria growth, but are still imprecise. Importantly, most statistical models are tuned to maximize overall accuracy (e.g., high *R*^2^), but elevated FIB levels are often rare events. Thus, models frequently have low sensitivity; that is, the ratio of accurately estimated elevated FIB levels over instances of actual exceedences.

### 2.2. Chicago Prior-day Nowcast Model

Chicago Park District measures water quality of 20 beaches along 26 miles against Lake Michigan on Chicago’s eastern shore shown in Figure 1. There is no single source of microbial contamination in Lake Michigan; rather it is likely introduced by birds, land-based animals, beaches, and people. The Chicago River is contiguous to Lake Michigan but a lock limits water flowing into the lake. While the lock is sometimes opened, it is generally less than once per year, and beaches are immediately closed (Chicago Metropolitan Water Reclamation District 2016).

Beaches operate from Memorial Day weekend, which is just before the last Monday in May, through Labor Day, which is the first Monday in September – approximately 122 days. As a result, there are approximately 2,440 “beach days,” which each represent an observation in the model. If FIB levels exceed the acceptable threshold for a given beach day, a public advisory notice is issued, warning potential swimmers about the heigtened risk.

Between 2011 and 2016, Chicago Park District placed hydrometeorological sensors to automatically collect covariates on water and atmospheric conditions. Buoys were installed at five Chicago beaches–Foster, Montrose, Oak, 63rd Street, and Calumet–to collect turbidity, wave height, wave period, water temperature, and depth of the sensor. Weather stations were installed at three beaches–Foster, Oak, and 63rd Street–and collected wind direction, wind speed, air temperature, rainfall, solar radiation, relative humidity, and barometric pressure. Data were collected from the hydrometeorological sensors between every 2 and 5 minutes from May through Setpember and aggregated.

Water samples were collected for culture-based testing each morning and recorded, usually around noon. Sampling was done on weekdays; however, weekend and holiday sampling was conducted if the prior readings were elevated or if Chicago Park District determined the weekend was busy. Between 2006 and 2016, only 3 percent of the samples were taken on weekends.

Shively et al. (2016) used this data to build a prior-day nowcast model which used the prior-day culture tests and same-day hydrometeorological data to predict FIB levels at all beaches. The novel model leveraged the sensors to automate the management process by providing estimates to beach managers each weekday.

Predictions were obtained from a random forest model and the predictions were published online for beach visitors. In 2015 and 2016, the overall accuracy was 90 and 93 percent, respectively, and specificity (true negative rate) was 98 and 99 percent. However, the sensitivity (true positive rate) was only 7 and 11.9 percent for all Chicago beaches.

Beginning in 2015, Chicago Park District began to use limited qPCR testing of *enterococci* at 5 beaches. Data were collected but not incorporated into the predictive model. During the summer of 2017, qPCR testing was expanded to all 20 beaches and culture-based testing of *E. coli* was paused.

### 2.3. Hybrid Nowcast Model

Whitman and Nevers (2008) observed that FIB levels at Chicago beaches often correlated with nearby beaches on the same day, where extreme highs and extreme lows were simultaneous for most beaches. Figure 2 shows the correlation of culture-based *E. coli* measurements between Chicago beaches from 2006 through 2017. The branches denote the “nearest neighbor” for each pair of beaches, a simple way to show similar beaches. Beaches were not uniformally correlated. Some beaches displayed strong clusters of correlation that exceed the correlation of readings within the same beach between days.

We identified clusters of beaches that had correlated FIB levels. A clustering algorithm, *J* (…), assigned beaches, *x_i_*, to a cluster *k* based on correlation between historical bacteria levels between 2006 and 2016:

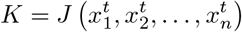

We selected one beach, 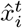, in each cluster to be the feature beach for predicting bacteria levels for the remaining *m* beaches in the cluster, 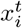. Thus, each cluster (*k*) had the following membership: 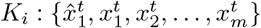.

To generate the predictions, we formulated a model that limits predictions to each cluster:

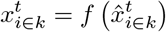

so the feature beach 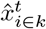, the *i*^th^ beach in cluster *k* used data from time *t* to predict the remaining beaches 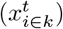 in the same cluster in the same time period.

This model leverages observations from the same time *t* to predict FIB levels at the other beaches on the same day. The rapid results from qPCR testing of *enterococci* allows recreational beach managers to observe beach readings at selected beaches, 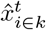, to predict culture-based *E. coli* levels at other beaches. The factors and conditions which led to elevated levels are captured and preserved in the model to render predictions.

### 2.4. Identifying Beach Clusters

We applied the above approach to Chicago’s beach data. We used K-means clustering algorithm to detect when beaches had similar movements. First, there was an initial, random guess for which beaches belong in clusters. The distance or error was then measured for all clusters. Then, membership of clusters was slightly altered, and we calculated whether the error had increased or decreased. This process was repeated until the lowest measured error was obtained.

Mathematically:

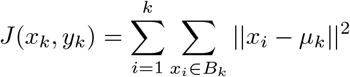

We chose to limit to 5 clusters (*k* = 5) because it aligned to the number of qPCR testing sites by Chicago Park District starting in 2015. K-means was applied to beaches based on latitude, longitude, total *E. coli* exceedances, and length of the longest breakwater. Each of these variables was scaled and centered by calculating z-scores prior to clustering.

Some beaches were removed from the clustering because they were historical outliers for bacteria levels or had distinct physical features. We removed beaches that have lengthy breakwaters since it has an impact on bacteria levels at those beaches. We tested this hypothesis by measuring the distance of the southernmost part of the beach to the northeasternmost edge of the breakwater. The correlation between the breakwater length and total bacteria exceedances between 2006 and 2017 was positive (*r* = 0.73). We removed 63rd Street, Rainbow, Montrose, and Ohio, which have long breakwaters or a similar feature.

Likewise, our earlier analytical modeling showed that beaches with a high frequency of high bacteria levels often confounded the model. Calumet, which had high exceedances as well as a medium-sized breakwater, was also removed from the analytical model. These 5 removed beaches comprised two of the clusters from the initial K-means analysis.

The final list of beaches and their respective clusters are listed in Table 1 and mapped in Figure 1.

**Table 1:** Final results of K-means clustering. The ^ denotes feature beaches used to predict remaining beaches in the cluster.

| $k$ | Cluster 1 | Cluster 2 | Cluster 3 | Cluster 4 | Cluster 5 |
| --- | --- | --- | --- | --- | --- |
| $\hat{x}$ | Foster$\hat{\cdot}$ | North Avenue$\hat{\cdot}$ | Leone$\hat{\cdot}$ | 31st$\hat{\cdot}$ | South Shore$\hat{\cdot}$ |
| $x_{1 \in k}$ | Osterman | Oak Street | Juneway | 12th | 57th |
| $x_{2 \in k}$ | Albion | | Howard | 39th | |
| $x_{3 \in k}$ | | | Rogers | | |
| $x_{4 \in k}$ | | | Jarvis | | |

Within each cluster, the beach with the most historical culture-based *E. coli* exceedances was selected to be the feature beach whose *enterococci* qPCR result would be input to the model. By rapid testing the beaches with the most frequest exceedances, we captured an added operational benefit of maximizing the number of correct advisories. The remaining beaches were selected to be predicted by the model.

### 2.5. Predictors and Covariates

Previous literature has focused on using a variety of covariates to predict *E. coli* levels. We created and tested an analytical model which relied on interbeach correlations and also used other variables cited in previous literature to predict FIB in Lake Michigan. The model resembled the model constructed in section 3 of the paper.

Ackerman and Weisberg (2003), Morrison et al. (2003) found that large rainstorms increased FIB levels 2 to 5 days after the rain has ceased. We collected data from the Dark Sky application programmer interface (API) to understand forecasted rainfall for a day of water measurement and also understand various historical measures of rainfall.

Likewise, the same data source was used to determine the amount of sunlight the lake was exposed to at the beaches. Whitman et al. (2004) found that increased sunlight inactivated *E. coli* at a Chicago beach, whereas cloudy days decreased *E. coli*. Similar to precipitation, Dark Sky data was used to measure forecasted cloud cover as well as historical measurements of cloud cover preceding water quality tests.

We included various measurements of wind since Smith et al. (1999) and Olyphant and Whitman (2004) found that wind is a reliable predictor of FIB levels. In particular, beaches downwind from point-pollution sources or where FIB was active are more likely to experience elevated *E. coli* levels. We used the Dark Sky API to determine the wind direction and intensity each day, as well has historical data preceeding a beach sample.

Similarly, wave and tidal patterns seem to influence observed FIB levels. Le Fevre and Lewis (2003) found that *E. coli* levels could be resuspended under tidal waves, allowing *E. coli* to be observed in locations only reached by waves or tidal levels. In a multivariate model, Crowther, Kay, and Wyer (2001) showed that tidal levels affected observed FIB levels. Unfortunately, our team did not have access to suffucient historical data to determine tidal levels, but we approximated their effect by using the lunar phase. Additionally, we used the previous day’s Lake Michigan water level and 3-day average water level, as measured and obtained by the National Oceanic and Atmospheric Administration (NOAA) at Calumet Harbor in Chicago.

Finally, researchers have found the human and animal activity at the beaches can influence FIB levels. Animal activity, whether domesticated pets or wildlife, may introduce fecal bacteria to the environment (Boehm et al. 2003; Reeves et al. 2004). Likewise, human activity alongside rain and runoff can introduce bacteria from the beaches into the water. While Chicago Park District does not have frequent surveys of beach visitors for each beach, we used indicators for weekday and weekend as well as the Julian date to approximate activity at each beach.

Table 2 shows the covariates considered.

**Table 2:** Variables used for multivariate Hybrid model

| Variables |
| --- |
| **Precipitation** |
| Forecast for today |
| Yesterday total rainfall |
| 3 day total rainfall |
| Total rainfall total until 8am |
| **Sunlight** |
| Daily cloud cover forecast |
| Prior-day cloudiness |
| 3-day total cloudiness |
| Length of daylight time in a day |

Table 2: Variables used for multivariate Hybrid model (*continued*)
| Variables |
| --- |
| **Wind** |
| Wind Direction |
| Wind Speed |
| 1-day average wind speed |
| 3-day average wind speed |
| Wind speed at 8am |
| **Tidal Levels** |
| Lunar phase |
| **Lake Level** |
| 1-day lake level |
| 3-day average lake level |
| **Visitor Density** |
| Indicator for weekday |
| Julian date |

### 2.6. Building the Predictive Model

A random forest regression model was trained with the predicted beach’s name and 10 years of culture-based *E. coli* test results for each of the feature beaches. Variable importance for the random forest model was assessed by calculating the percentage increase in mean squared error (MSE) as a result of permuting the values for each variable (Figure 3). The most important feature was the *E. coli* level at Leone Beach. In fact, the importance of each beach’s *E. coli* level decreased as the beach location was further south. This is likely due to the fact that more beaches on the north side (10) are being predicted than on the south side (5) and due to the north-to-south current of the lake.

The name of the beach being predicted had the lowest importance. Without this predictor, the predicted *E. coli* for a specific day would be the same for each beach. Indeed, because of the correlation that exists between certain beaches, it is not surprising that beach identity would rank last against the importance of *E. coli* levels along the whole lakefront. Still, permuting the beach name variable led to a 15.6 percent increase in the MSE.

The random forest regression model predicted *E. coli* levels in CFU/100ml units. These raw levels were transformed to a binary advisory decision. Due to left censoring at 1 CFU/100ml and an abundance of low *E. coli* readings, a limit can be observed at the lower left portion of a plot of residuals and fitted values.

As the number of trees in the random forest model grew, model error decreased until around 400 trees. No performance benefit was observed beyond 400 trees.

### 2.7. Validation

The model was tuned and validated using leave-one-year-out cross validation, in which the validation set consisted of all the observations in one year, and the training set consisted of all other observations. The process was repeated until each year had been in the validation set. The year 2016 was left out as a final validation set.

Historically, the Chicago prior-day model from 2015 through 2016 had a false-positive rate of 1.8 percent for all beaches. That is, the model falsely predicted elevated bacteria levels 1.8 percent of all beach days over the course of the summer–typically 30 beach-days per season. We chose to align the parameters of our model to this historical false-positive rate. As Figure 4 shows, we could have increased the number of times we correctly predicted elevated FIB levels, but it would come at a cost of also increasing the number unnecessary warnings to beach-goers.

Figure 4 shows the receiver operating characteristic (ROC), which shows the trade-off between increasing the number of days with true predictions versus days with falsely identified elevated levels. By constraining the model to only have a 1.8 percent false-positive rate, the true positive rate is forecasted to be 22 percent.

A threshold was chosen to transform the model prediction to a binary outcome. To keep the model’s false positive rate (FPR) near 1.8 percent, the threshold within each year that corresponded to a 1.8 percent was noted, and the mean threshold was then used to generate predictions for the holdout validation set.

## 3. Results and Discussion

To correctly assess the accuracy of the model, we needed to use out-of-sample predictions. As can happen with machine learning, it is possible that the accuracy of the model was caused by the analysts tuning the model to fit existing data and accompanying noise. Fitting existing data too well can lead to the model failing to accurately predict when encountering new data. Pilot analytical models were created using historical data from 2015 and 2016, using the modeling concept described in Section 2. Pilot models were live tested at Chicago beaches in 2017 on newly collected data.

In summer 2017, the pilot analytical model went live with some limitations imposed by available data. The model inputs were *enterococci* qPCR test results for Calumet, Rainbow, South Shore, 63rd Street, and Montrose, and the predicted beaches were the other 15 regularly tested beaches. The model was trained using results from the qPCR pilot for 2015 and 2016, and predicted the culture-based *E. coli* levels for the predicted beach on the same day. An application was developed and deployed which regularly checked Chicago’s public data portal for qPCR results for the feature beaches. Every day during the summer, once all the qPCR results were posted, the application automatically ran the model and uploaded the predictions on the public data portal within five minutes.

Beginning on Friday, May 26, water samples were collected by the Chicago Park District and qPCR testing was completed by the University of Illinois at Chicago. Those results were then posted to the City of Chicago Data Portal–typically around noon (City of Chicago 2016b). About 5 minutes later, we posted the predictions based on our model to the Data Portal (City of Chicago 2016a). The last predictions and samples were conducted on September 4th. While qPCR data were available for all beaches, we ignored any qPCR reading except for the 5 beaches used as model inputs.

Both predictions and samples were collected each day during the week. This process was different from prior years where samples were usually collected only during weekdays and non-holidays, except when levels had been elevated on the previous Friday or it was suspected to be a busy weekend on the beaches.

To translate the predicted *E. coli* level to an advisory decision, beaches that were forecasted to exceed 381 CFU/100ml were predicted to have a swim advisory due to FIB. Even though this threshold exceeds Environmental Protection Agency (EPA) requirements, that threshold was empircally shown to be the best correlate to actual exceedances using EPA standards. Subsequent lab testing and swim advisories due to FIB were issued only if FIB exceeded EPA standards.

Over the summer of 2017, we compared predictions to the results from the actual qPCR *enterococci* test results (City of Chicago 2016c) and calculated several standard measurements to evaluate the performance of the model. We specifically compared the predictions at the 15 beaches where the hybrid modeling was used to formulate predictions, not on all 20 beaches because some were not predicted, but were inputs for predictions at other beaches.

### 3.1. Results

Table 3 shows the hybrid analytical model had a sensitivity (true positive rate) of 11.2 percent for the predicted beaches compared to an average of 3.4 percent at the same beaches in 2015 and 2016. That is, the hybrid model increased the true positive rate by 230 percent over the historical average. Additionally, these beaches have typically not been predicted as well as the other beaches – the same prior model that averaged 3.4 percent for the 15 predicted beaches averaged 9.5 percent for all 20 beaches. The precision of the model–the portion of predicted high FIB that were accurate compared to all predicted FIB beach days–also grew from 17 to 27 percent–a 59 percent increase. However, the false positive rate was slightly higher at 2 percent during 2017 compared to our 1.8 percent target.

**Table 3:** Comparing Specificity, Sensitivity, Accuracy, Area Under Curve (AUC), and Matthews Correlation Coefficient (MCC) between Hybrid, Multivariate, and Prior-day Nowcast models for the 15 pilot beaches.

| Model | Specificity | Sensitivity | Precision | Accuracy | AUC | MCC |
| --- | --- | --- | --- | --- | --- | --- |
| 2017 Hybrid | 0.980 | 0.112 | 0.273 | 0.926 | 0.753 | 0.142 |
| 2017 Multivariate | 0.971 | 0.060 | 0.125 | 0.912 | 0.738 | 0.044 |
| 2016 Prior-day | 0.988 | 0.037 | 0.167 | 0.930 | 0.644 | 0.052 |
| 2015 Prior-day | 0.984 | 0.032 | 0.176 | 0.892 | 0.655 | 0.036 |

Area Under Curve (AUC) is a measure that shows the trade-off between the true positive rate and false positive rate for the model. The AUC for the hybrid model is 0.753, which is higher than the prior-day model that was around 0.64 in 2015 and 2016. A higher AUC does indicate the model has improved the ability to estimate beaches with elevated bacteria levels without raising a disproportionate number of false positives.

The Matthews Correlation Coefficient (MCC) provides an approximation for the goodness-of-fit for predictors. Like other correlation coefficients, it ranges between +1 (perfect prediction of elevated FIB levels) to −1 (complete disagreement). Unlike AUC, it accounts for all four classification outcomes: true positives, true negatives, false positives, and false negatives. The MCC for the hybrid model was substantially higher at 0.142 compared to 0.052 and 0.036, respectively, for 2016 and 2015. It is another indication in the relatively improved model performance compared to prior-day modeling.

We also compared performance to a model that combines interbeach correlation with predictors for precipitation, sunlight, wind, proxies for human density, and tidal and lake levels discussed in section 2.6. The performance of the multivariate hybrid model was similar to the hybrid model without other predictors. AUC for the multivate hybrid model was 0.738, 0.015 lower. The MCC for the multivarite model was substantially lower for the multivariate model at 0.044. Key metrics for sensitivity, 6 percent, and precision, 12 percent, were lower than the hybrid-only model. This indicated that our predictors only provided marginal improvements to the precision of the predictions, where most of the predictive power is provided by the interbeach correlation.

Accuracy of the hybrid nowcast model was in line with the previous performance of the Chicago prior-day nowcast model. Since most days do not have elevated *E. coli* levels, the increase in sensitivity (true positive rate) did not make a major impact on overall accuracy, which remained around 93 percent.

### 3.2. Discussion

Attempts in prior literature to forecast FIB levels in beaches have used essentially the same functional form (Nevers and Whitman 2005; Frick, Ge, and Zepp 2008; Hou, Rabinovici, and Boehm 2006). Lab data on FIB from the previous day are combined with hydrometeorological and other predictors to predict whether FIB levels will exceed the suggested thresholds (Ackerman and Weisberg 2003; Whitman et al. 2004; Smith et al. 1999). Recent innovations, such as hydrometeorological sensors (Shively et al. 2016), have enabled novel ways to collect the predictors leading to improved accuracy and reduced latency. Likewise, more sophisticated algorithms, such as machine learning and genetic algorithms (Cyterski et al. 2014; Brooks et al. 2016), have been used to improve performance by allowing for more complex interactions between predictors. Yet, the concept of these models still predominately relies on prior-day laboratory results, which we’ve dubbed the “prior-day nowcast model”. Evidence suggests that the contributors to high FIB levels do not persist from day-to-day (Morrison et al. 2003; Cheung, Chang, and Hung 1991; Brenniman, Rosenberg, and Northrop 1981), which can explain why many attempts to predict FIB levels in beaches have not sustained high levels of accuracy.

Previous research has found that FIB levels in Chicago’s beaches are highly correlated and rarely encounter consecutive days of elevated FIB levels Whitman and Nevers (2008). At the same time, qPCR testing has become more widely used, but is still expensive (Bienkowski 2017). Because qPCR testing provides immediate results, we proposed the hybrid nowcast model to use limited qPCR data from selected beaches to predict FIB levels in other beaches by exploiting correlations between beaches. The hybrid nowcast model removes the dependency on day-old FIB information commonly used in other models. This approach more closely resembles a “missing data” problem, where we are attempting to “fill-in” the missing values (beaches without qPCR testing). For beach networks that are highly correlated, like Chicago’s, hybrid nowcasting was able to increase model sensitivity 230 percent without increasing the rate of false positives. When a beach was predicted to have an elevated level, it was 59.2 percent less likely to have been a “false positive” (type I error). The increase in correct predictions, however, was not induced by simply raising more erroneous warnings (type II errors) as those only increased two-tenths of a percent.

In terms of impact on beachgoers, the hybrid model resulted in 90 correct advisories while there were 71 incorrect advisories. In 2016, the Chicago prior-day nowcast model issued 12 correct advisories while issuing 184 incorrect advisories. Likewise in 2015, the prior-day model issued 14 correct advisories versus 184 incorrect advisories. The difference in performance was likely due to the use of qPCR to obtain FIB levels on the same day. Since Chicago beaches rarely encounter consecutive days of elevated FIB levels, prior-day readings do not accurately predict elevated levels on the next day. The factors that cause FIB levels to grow rarely persist between days as few beaches have consecutive days of beach warnings. However, elevated FIB levels portend ideal conditions for FIB in other beaches on the same day.

We explored the addition of other predictors and covariates, but it did not increase the model performance over the model only using interbeach correlation. Information and predictive-power provided by covariates have already been captured by the qPCR testing conducted at other beaches. For instance, while rainfall can approximate the conditions for FIB growth, the variation of FIB levels at other beaches is a better predictor since it is reliant on actual beach conditions. The temporal linkage between observations for ideal FIB conditions and actual FIB growth has been hard to demonstrate in a large ecosystem like Lake Michigan. Meanwhile, the interbeach correlations seem to provide a better signal that ideal conditions already exist.

Nothwithstanding, we also realize our hydrometeorological data is based on third-party sources instead of active sensors in the water which may be more accurate and stronger predictors. Chicago beaches do have hydrometeorological sensors in place, but those data are only available for a couple of years. Our measurements for visitor density are very loose approximations based on available data, but does not mirror the methodology used in studies that have shown the relationship between human activity and FIB levels. These could be improved in the future and may provide better capability for prediction.

Although Chicago was able to deploy qPCR testing at all 20 beaches, the cost and complexity of qPCR equipment limits widescale deployment. Moreover, we removed four beaches from this model because of physical characteristics, such as long breakwaters, that prevented them from being clustered with other beaches; one beach was removed due to frequently high readings; and we used 5 beaches to predict culture-based *E. coli* levels. As a result, 10 total beaches would need to undergo *enterococci* qPCR testing and 10 beaches would be able to forgo any testing. Of the 10 undergoing qPCR testing, 5 would be tested to estimate FIB levels for 10 other beaches and the other 5 would need qPCR testing since they have distinct geographies, breakwaters, or other unique physical characteristics.

The hybrid model used predictor beaches that were more prone to elevated FIB levels than all other beaches in each cluster: Foster, North Avenue, Leone, 31st, and South Shore. Moreover, we removed Calumet from the model since it frequently had elevated levels. Beach monitors could levarage a similar strategy of deploying qPCR testing to frequently contaminated beaches and relying on statistical models for the remaining beaches. This will naturally improve model performance since beaches which comprise the mode number of issues will be accurately measured, leading to more accurate beach warnings, and limits expensive equipment needed for rapid testing.

While this paper showed a 230 percent increase in sensitivity, the ability to predict elevated FIB levels is still relatively low. One method to improve sensitivity without increasing AUC is increasing the rate of false positives. This method certainly would be an inconvenience for beachgoers, but also increase the chance of warning beachgoers of elevated FIB. However, there are a lack of guidelines or studies on acceptable levels of false positives. Our target rate was determined based on the average false positive rate for prior years. It may be worthwhile to incorrectly predict additional beach warnings if it deters exposure to FIB, especially for children or those with weak immune systems. Research on predicting elevated FIB can be guided by further research on the economic impact of incorrectly providing warnings when none were warranted. That analysis can help this strand of research determine a target for an appropriate level of false negatives.

## 4. Conclusions

- Hybrid nowcast modeling preserves information from the current day, instead of test data from the prior day.
- The modeling technique allows for limited deployments of the costly qPCR testing equipment.
- Sensitivity of predicting elevated FIB levels increased from 3.4 percent to 11.2 percent–a 230 percent increase–while reducing false positive rates by 59.2 percent and maintaining similar rate for false negatives.

## 5. Acknowledgements

We are indebted to Chicago Park District staff, especially Cathy Breitenbach and Carol Kim, who provided incredible guidance and feedback while taking time to work with the research team. Meredith Nevers graciously provided the research team with her considerable expertise. We are grateful for Chi Hack Night which provided a forum for the volunteers to contribute to this project, in particular, we thank Forest Gregg who helped spur this project. We appreciate Jonathan Levy’s work to publish the necessary data to make it available to the research team and the public and Sean Thornton’s helpful comments and edits on early drafts. The University of Illinois at Chicago’s School of Public Health diligently completed all water testing in a timely matter. Finally, we wish to thank the attendees of State of Lake Michigan conference in 2017 for their comments and feedback.

This research did not receive any specific grants from funding agencies in the public, commercial, or not-for-profit sectors.

## Appendix A. Supplementary Materials

Raw data from beach reading and historical forecasts generated by the hybrid nowcast model and past swim advisories are available on the City of Chicago Data Portal (City of Chicago 2016b; City of Chicago 2016c; City of Chicago 2016a) and updated during the beach season. The statistical model developed and described in this paper is also available online and is open source (Lucius et al. 2018). Finally, the code used to develop this paper is entirely reproducable, meaning all code and formula to generate figures and tables are reviewable.

**Figure A.1:**
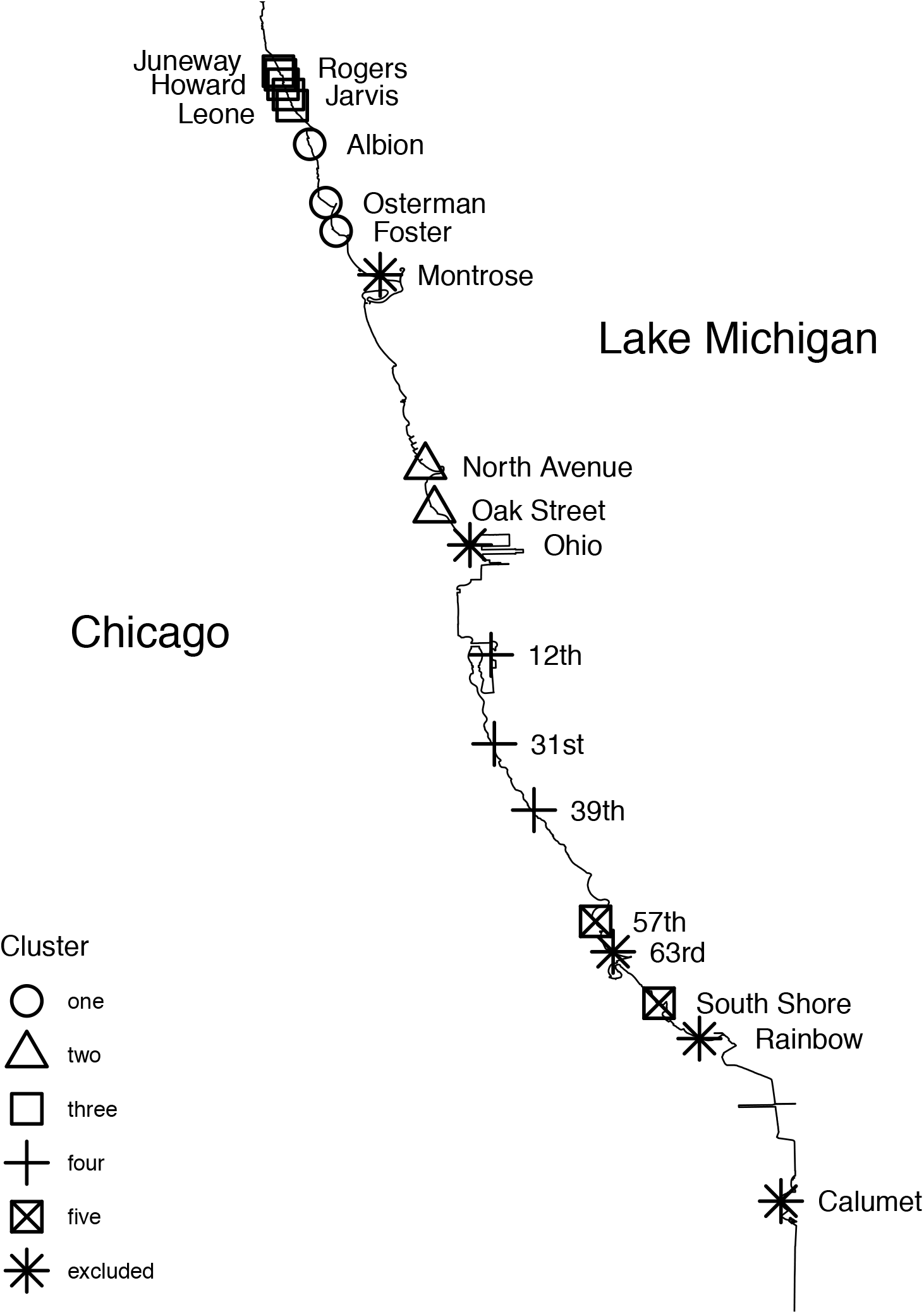
Map of Lake Michigan beaches in Chicago with results of k-means clustering.

**Figure A.2:**
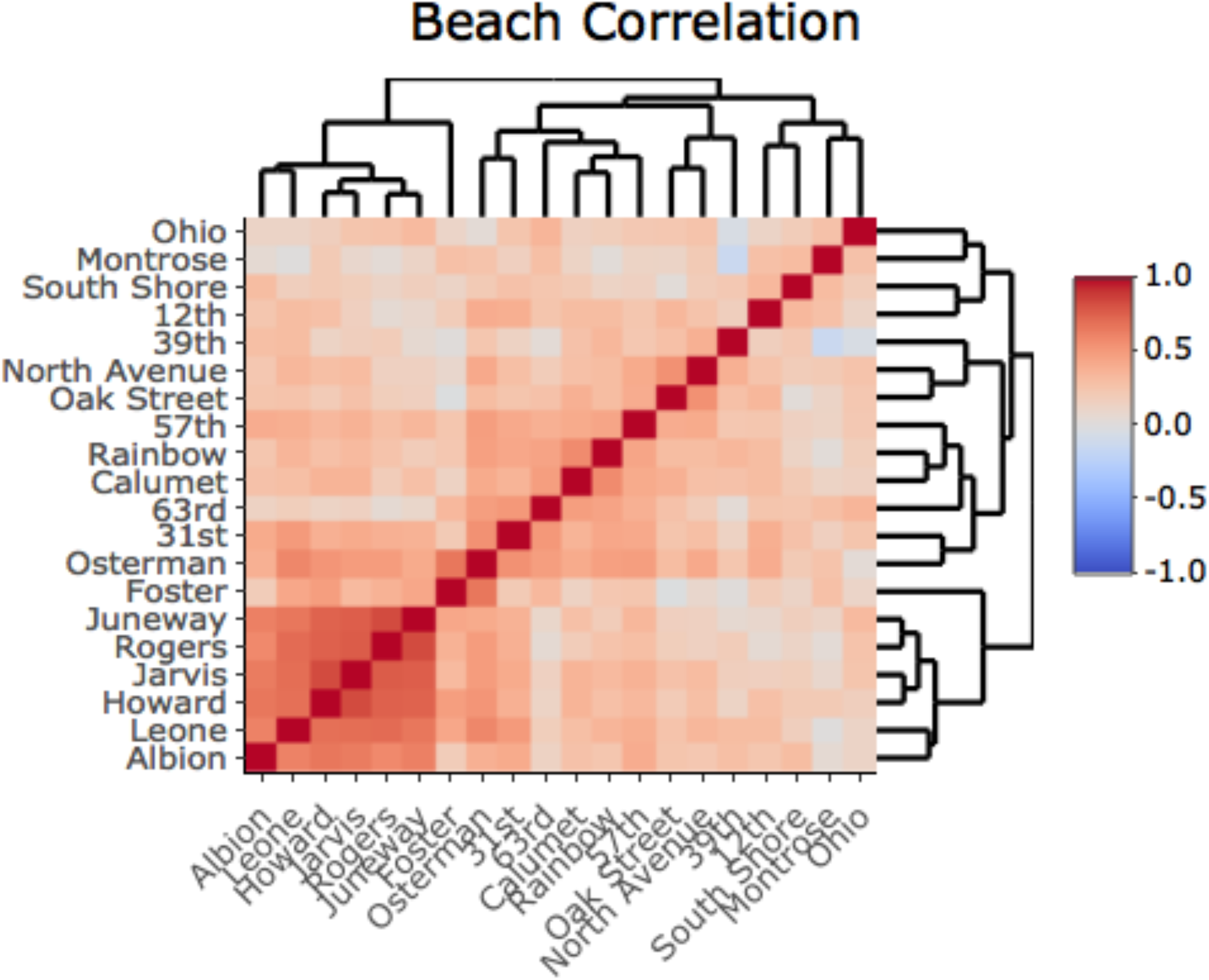
Pearson correlation coefficient heat map of daily E. coli levels at Chicago beaches between 2006 and 2017. Each level of tree shows the nearest-neighbor correlation.

**Figure A.3:**
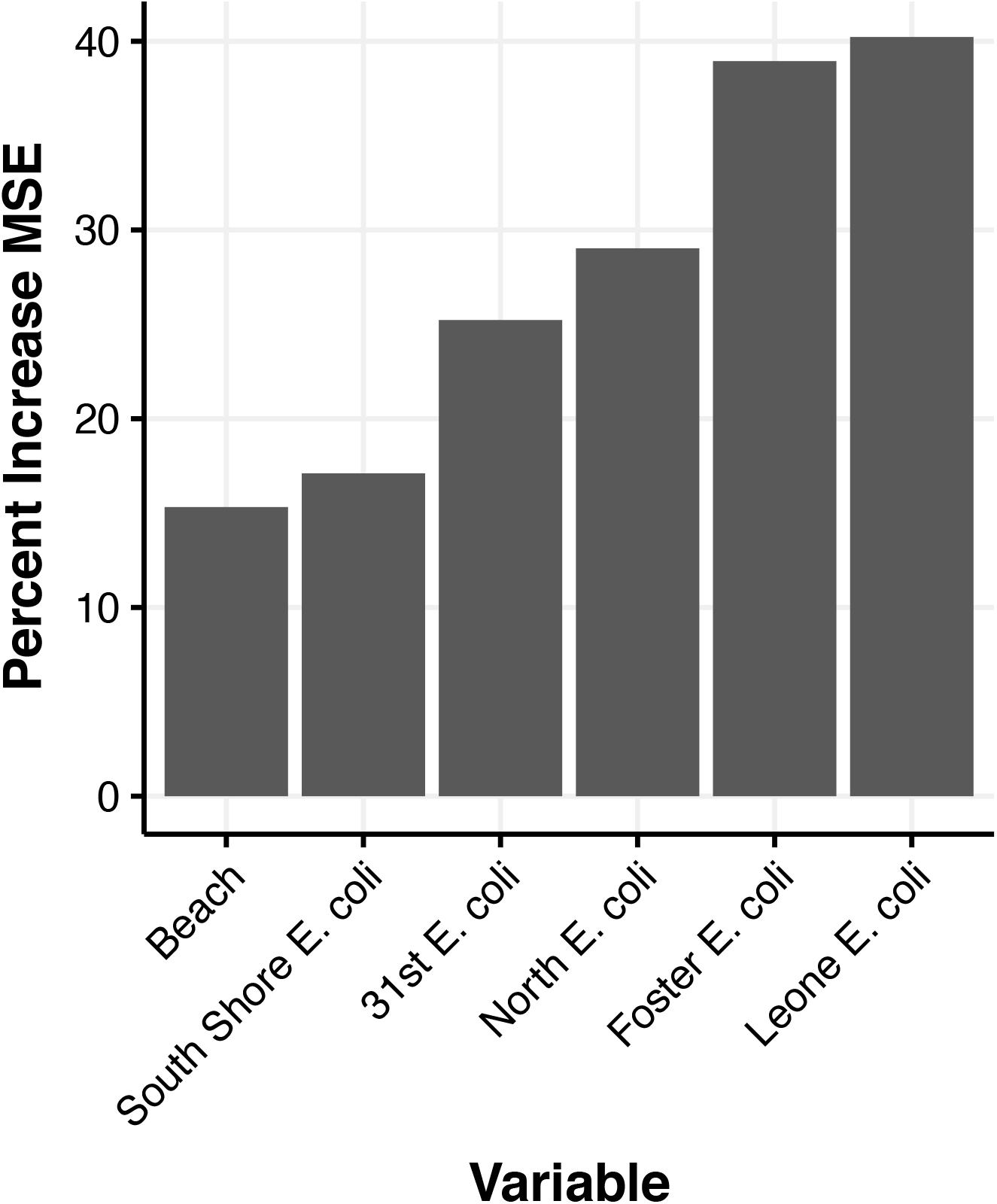
Variable importance of each factor when added to random forest model as measured by mean squared error

**Figure A.4:**
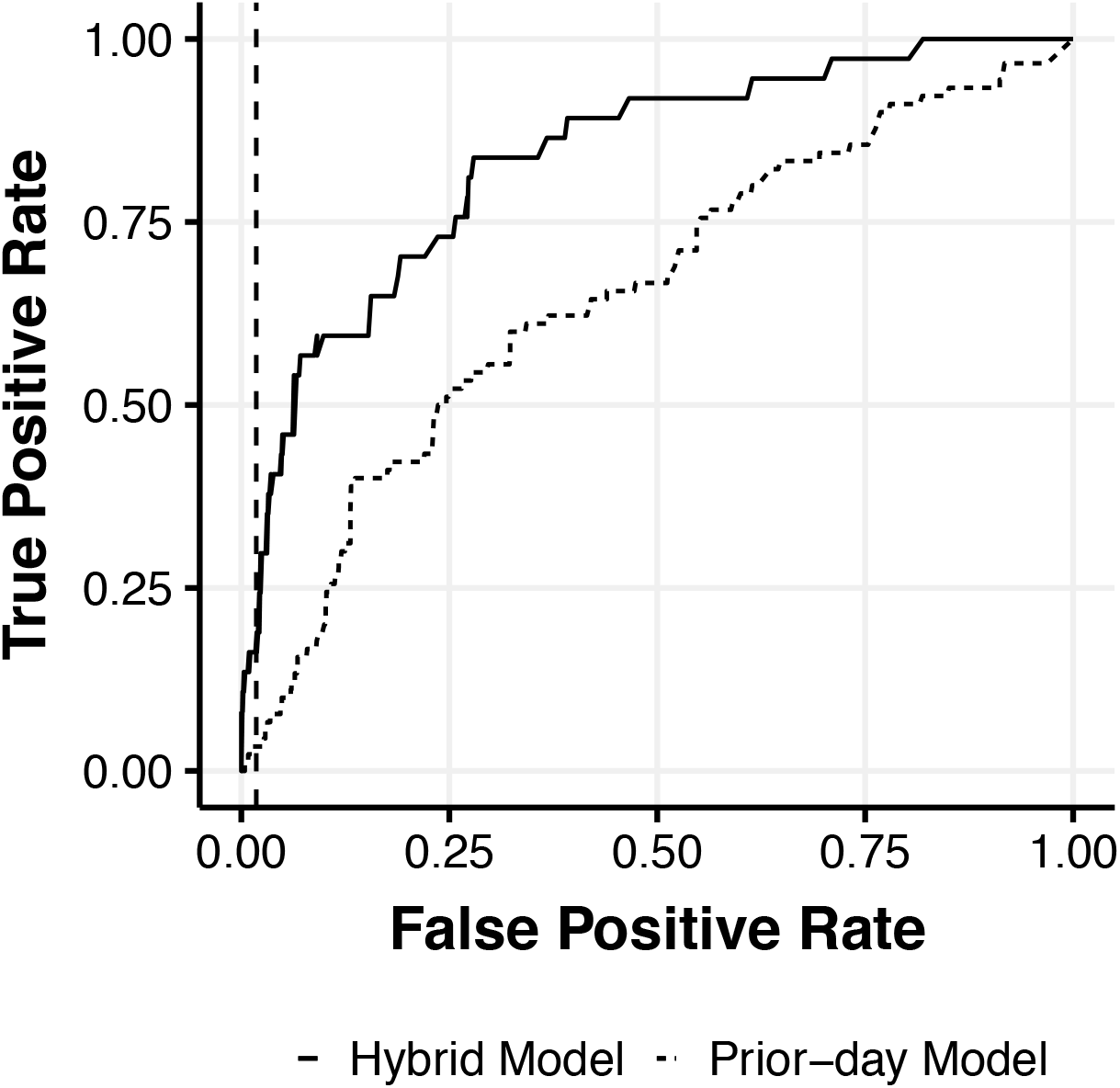
Receiver Operating Characteristic (ROC) for Hybrid Model and Prior-day Model. Dashed line shows historical false positive rate for prior-day model in Chicago.

**Figure A.5:**
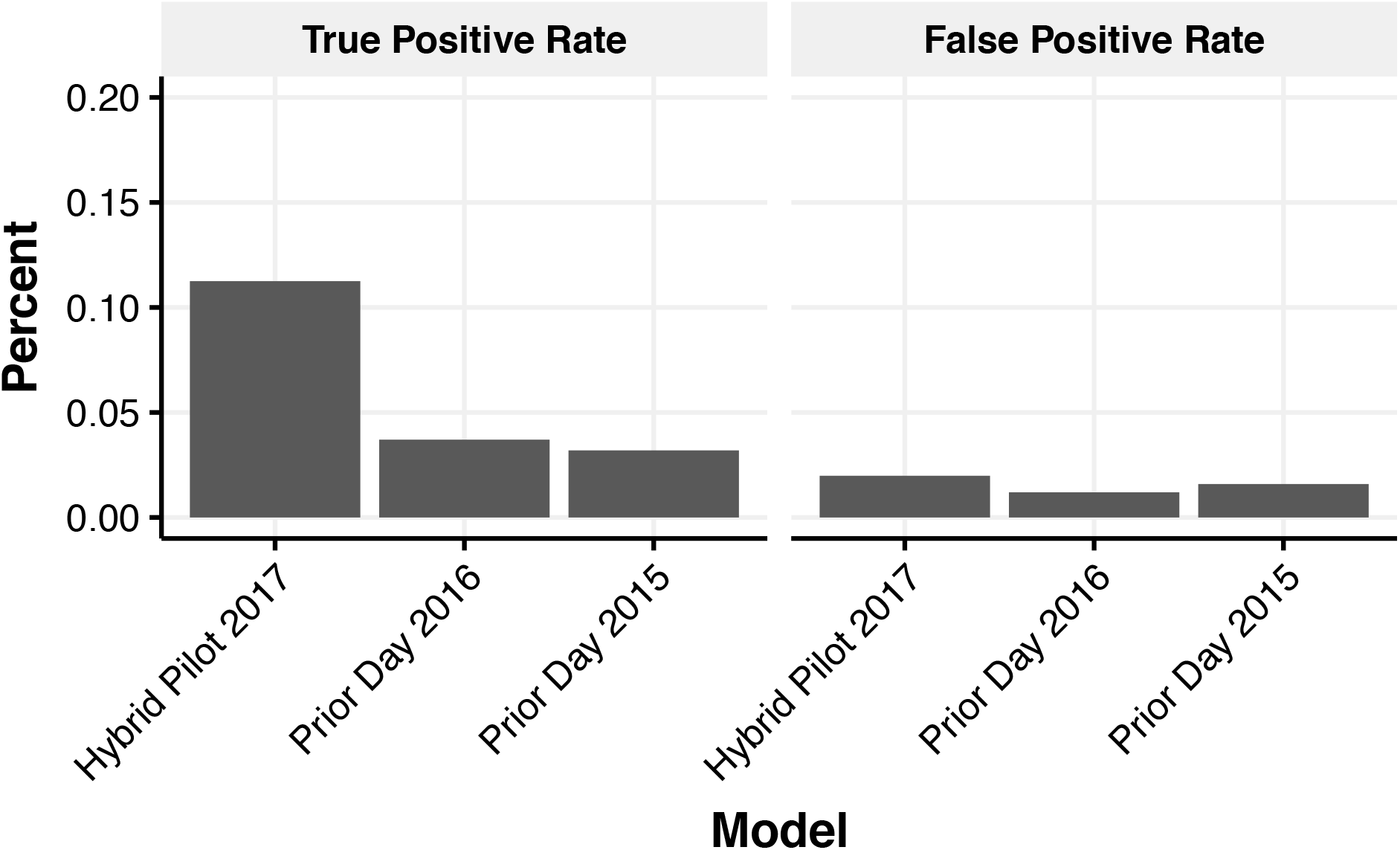
Comparison of the 2017 hybrid pilot to existing prior-day model for "true positive rates" (sensitivity) and "false positive rates" (type I error)

**Figure A.6:**
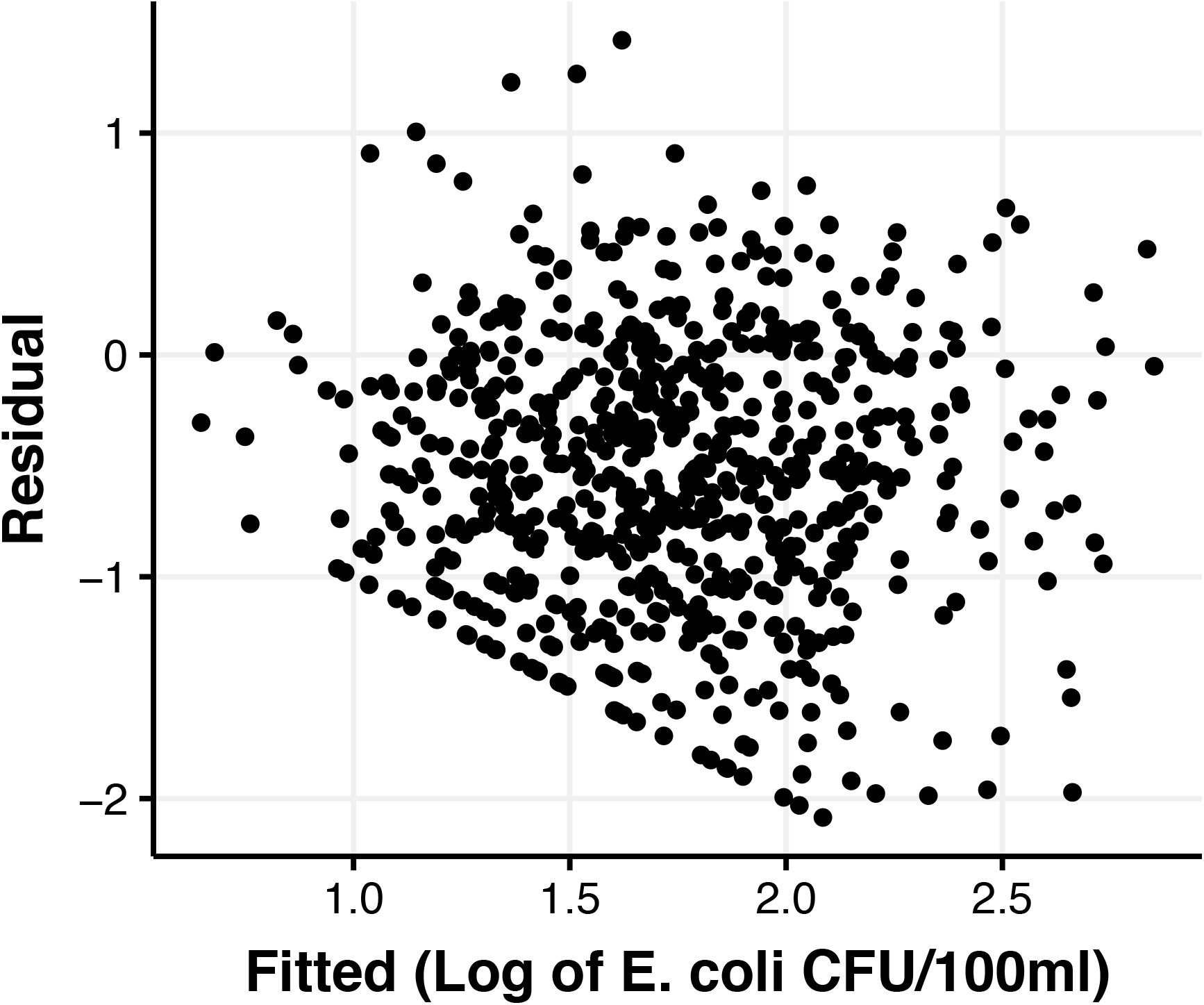
Plot of the log of raw fitted values versus residuals from the random forest model.

**Figure A.7:**
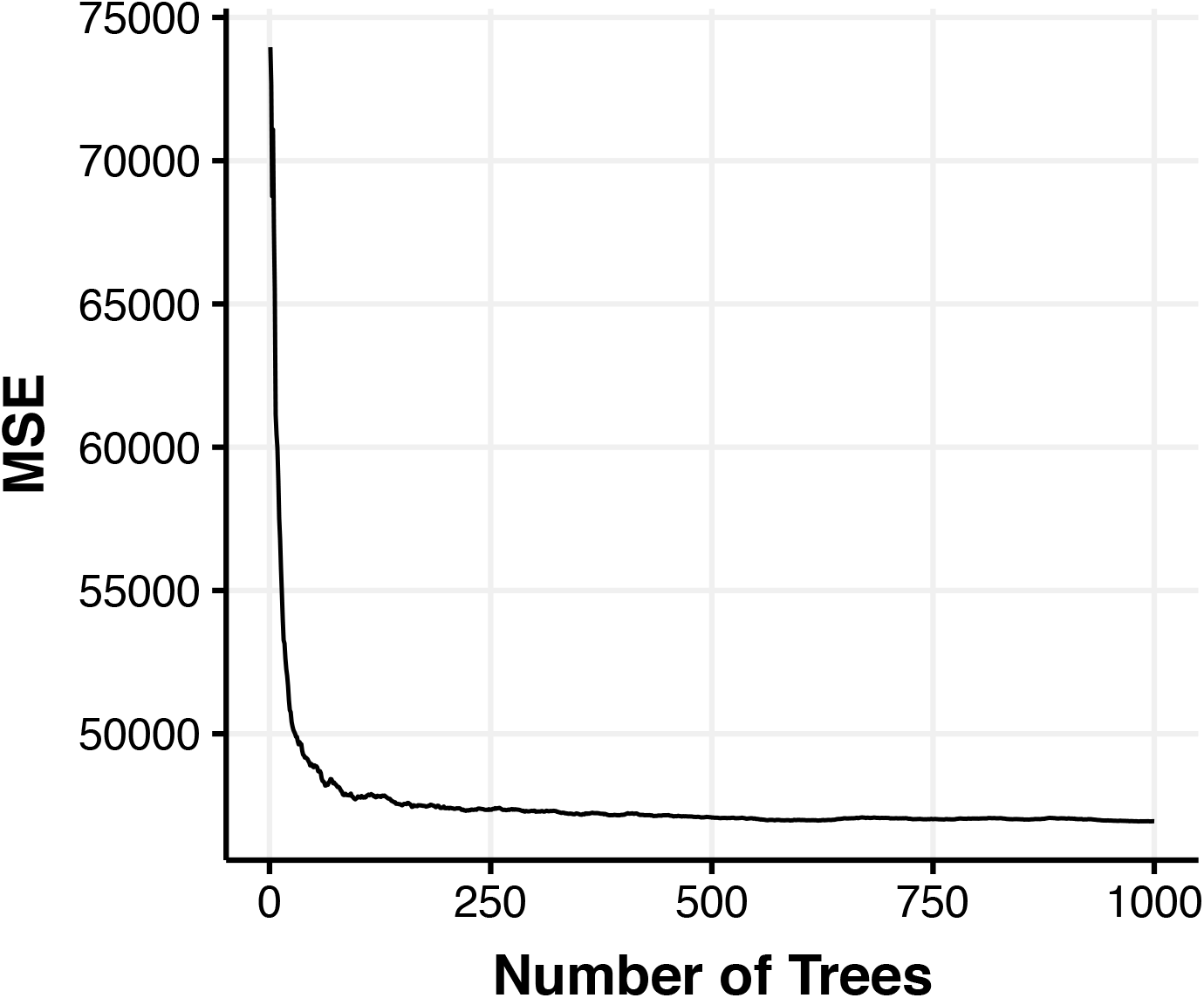
Model performance measured by mean squared error (MSR) as the number of trees grow within the random forest model.

